# Cold Vacuum Extracts of Double Cherry Blossom (Gosen-Sakura) Leaves Show Antitumor Activity

**DOI:** 10.1101/551424

**Authors:** Junko Shibato, Fumiko Takenoya, Takahiro Hirabayashi, Ai Kimura, Yusuke Iwasaki, Yoko Toyota, Motohide Hori, Shigeru Tamogami, Randeep Rakwal, Seiji Shioda

## Abstract

The present research examines the possibility of finding bio-molecular compounds from the double cherry blossom (termed as ‘Gosen-Sakura’ of Gosen-city, Niigata-prefecture, Japan) leaves, which have been long used in the preparation of the traditional Japanese sweet (wagashi) – ‘sakura-mochi’. Based on its indicated anti-microbial properties historically, our study provides a new low temperature vacuum extraction method for extracting ‘near natural form of water soluble leaf (cell) extracts from the Gosen-Sakura, and demonstrates the presence of some ‘novel’ compound(s) with anti-tumor cell lines proliferation inhibitory affects through the MTT assay. To our knowledge, no reports exist on the sakura tree ‘leaf (cell) extracts’ inhibiting tumor cell line growth. We further examined and compared the effects of known compounds with anti-tumor activity, coumarin and benzyl alcohol with Gosen-Sakura leaf extract; results lead us to hypothesize that the Gosen-Sakura leaf extract contains substance(s) other than the above 2 known compounds, with antitumor effect. Additionally, we speculate on the underlying mechanism of action of the Gosen-Sakura leaf extract by targeting cell division at the point of DNA synthesis and causing apoptosis. In conclusion, we present scientific evidence on the presence of a certain ‘novel’ biomolecule(s), with anti-tumor activity, in the Gosen-Sakura leaf which has been long used as a Japanese – the ‘sakura-mochi’.

## Introduction

Sakura (cherry blossom) is a generic term for deciduous trees of the family Rosaceae (sub-family, Amygdaloideae; Tribus, Amygdaleae; Genus, *Prunus*; Sub-genus, *P.* subg. *Cerasus*; Sectio, *P*. sect. *Cerasus*; Nothospecies, *Prunus* × *yedoensis*; Cultivar, *P. y*. ‘Jindai-akebono – *P. y*. ‘Somei-yoshino’) represented by *Prunus cerasus* or *Prunus yedoensis* [1–3]; and is very familiar to the Japanese with a long cultural history. Cherry blossoms are popular as landscapes and are planted everywhere in Japan appreciated not only for their natural beauty (sakura viewing) but also in food as ingredients in tea and Japanese sweets mostly for its fragrance following the salt fermentation of the petals. Other than the petals, the leaves of the sakura tree are also used in the preparation of Japanese sweets since ancient times, and famously called the ‘sakura-mochi’ [4]; the sweet is eaten all over Japan during the sakura season. The sakura-mochi is prepared from pounded glutenous rice which contains sweet red bean paste wrapped around by a ‘salted’ sakura leaf of usually the double cherry blossom [4]. Though there are no direct evidences to the said anti-microbial and anti-bacterial properties of the sakura-leaves, few researches on the antioxidant activity of *Prunus* species are available [5–8].

The reason behind the use of the leaves of the sakura tree, in particular the double cherry blossom leaves, for edible use is thought to lie in its numerous metabolites, including the presence of isoroflavone purnetin, its glycoside plunnetrin, flavonoid isoquercitrin, and merylose side of coumarin glycoside; and there are some related literature available from flowers and leaves [7,9,10]. These compounds have been reported to have sedation, antitussive, expectorant, insomnia, mental stability, and relaxing effects. Since ancient times, the boiled cherry leaves extracts liquid has been used for preventing skin roughness [11], has anti-inflammatory action for the skin [12], and suppression of melanin production [13,14].

Although the double cherry blossom is present all over Japan, we have focused our attention on a cherry tree variety (*Primus lannesiana* Wils. cv. Sekiyama) grown in the Miura-village park of Gosen-city [15; https://www.city.gosen.lg.jp/hyande/2/4/2281.html], Niigata-prefecture (North-West Japan) for the present study. The cv. Sekiyama is a representative variety of long-known Satozakura, which has been well planted after the Somei-yoshino cherry tree. This is a highly resilient double cherry tree, with a beautiful big flower in deep red, and thus it is also preferred in foreign countries being widely planted in the United States and the United Kingdom, Australia and elsewhere. In Japan, ‘Sekiyama’ is cultivated not only for viewing the blossoms during the sakura season but also for edible use of both flower (petals) and leaves, so it is highly popular.

Moreover, our research group is involved in exploring ways to effectively utilize diverse agricultural products from different parts of Japan [16] and the double cherry blossom (sakura cv. Sekiyama) was identified as a possible source of novel natural products for medicinal purposes, which is based on the recorded use of Sakura blossoms and leaves in Japanese history. For this purpose we have been utilizing the power of low-temperature based vacuum extraction method, analyzing ingredients of powder residue and their effective utilization therein [16]. Unlike the conventional steam distillation method, the low-temperature vacuum extraction method extracts under the conditions of low temperature and vacuum of up to 30 to 40 degrees to obtain the original natural scent and almost 100% aroma contained in the raw material without damaging the ingredients. This method results in obtaining the oil and cell extract and dry cell powder [16,17]. In simple terms the extract in liquid form, from the cell, is termed as the ‘cell-extract’. Since active ingredients such as polyphenols which cannot be extracted by steam distillation method are detected from the cell extract, which we believe can be a rich source of compounds not only for aroma and beauty, but also as a raw material for various uses in food processing using efficient solid-liquid separation. This also serves as a zero emission technology for the 21^st^ century.

In the course of research on utilization of this low-temperature vacuum extraction method, we found that the Gosen-city double cherry blossom (*Primus lannesiana* Wils. cv. Sekiyama; hereafter, referred to as Gosen-Sakura) leaf extract has antitumor effect, and which is presented in this study. As the cherry leaves contain coumarin, which is known to have antitumor effect [18,19], we also compared the cell extract with coumarin and other known compounds such as benzyl alcohol for anti-tumor affects. Further, the research looked into the mechanism of action of tumor effects and possible explanation for the same.

## Materials and methods

### Preparation of Gosen-Sakura leaf extract and cell dry powder

The leaf (cell) extract and cell dry powder were obtained from the leaves of Gosen-Sakura, harvested in Gosen-city (Niigata prefecture) [15] during 2015 and 2016 following the low-temperature vacuum extraction method as shown in Fig 1 (adapted from [17]). The extraction was done using the low-temperature vacuum extractor (FED-50-300, F.E.C. Co., Ltd., Aioi, Japan; [16]). The obtained cell extract was used for the following experiments. As required, the cell dry powder was also used for experiments.

**Figure 1.**
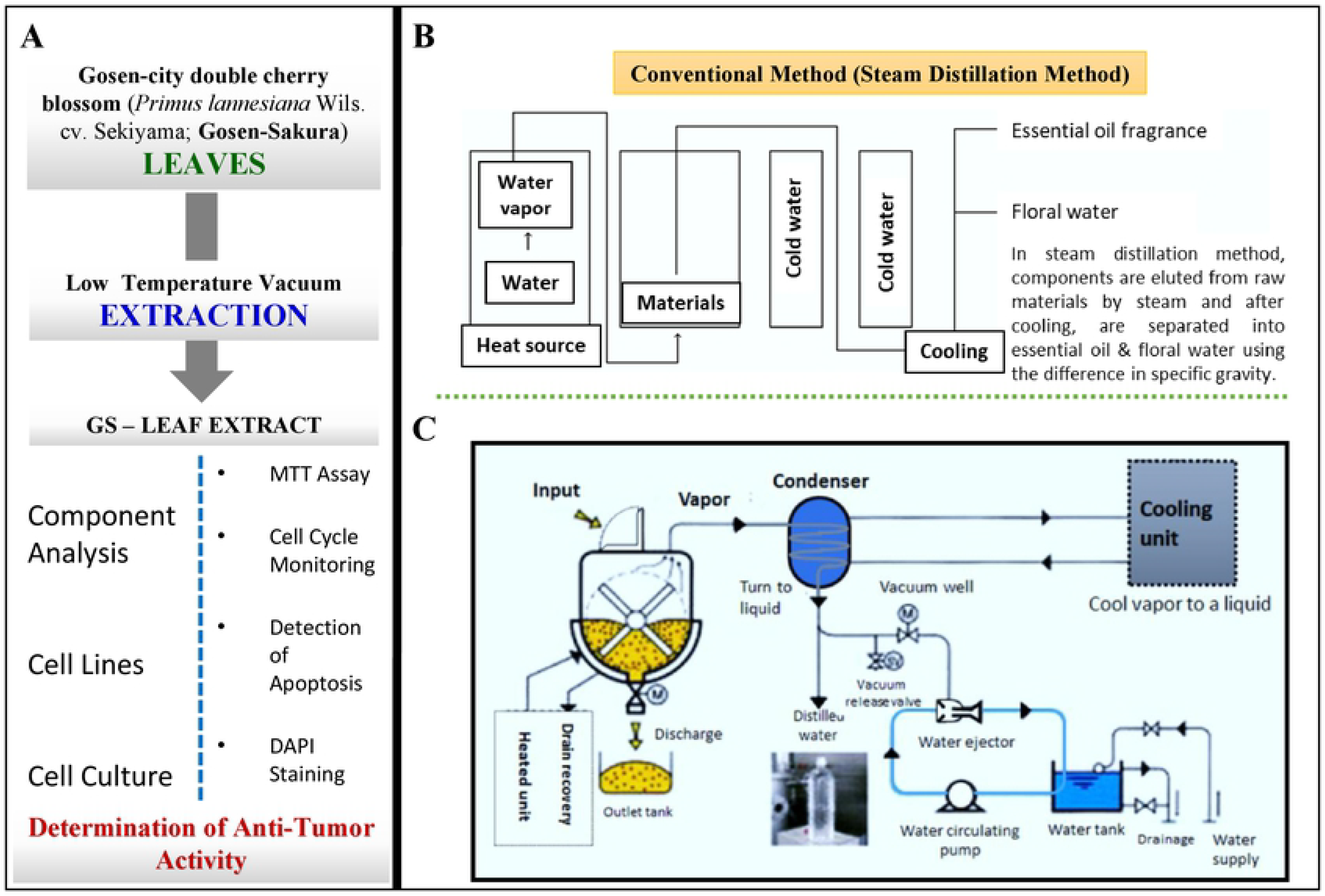
The Gosen-Sakura leaf extraction was carried out as per the standardized (and patented) protocol as described in Materials and Methods. A) The workflow, B) The conventional steam distillation method, and C) The schematics of the low-temperature vacuum extraction process equipment process and design. GS: Gosen-Sakura.

### Cell line

Cell lines HeLa (Human cervix epitheloid carcinoma: EC93021013), A549 (Human Caucasian lung carcinoma: EC86012804) and A431 (Human squamous carcinoma: EC85090402) were purchased from the ECACC (European Collection of Cell Cultures) collections (KAC Co. Ltd., Amagasaki-city, Japan). Cell lines HFSKF-II (Normal human fetal skin fibroblast: RCB0698), CACO-2 (Human colon carcinoma: RCB0988), HEK293 (Human embryonic kidney: RCB1637), HL60 (Human promyelocytic leukemia: RCB0041), HSC-3 (Human oral squamous carcinoma: RCB1975), LNCap.FGC (Human Prostate Carcinoma: RCB2144), MCF7 (Human cell line breast adenocarcinoma: RCB1904), HGC-27 (Human Stomach Carcinoma: RCB0500), OCUB-F (Human breast cancer: RCB0882), EC-GI-10 (Human esophageal cancer: RCB0774), Hep G2 (Human hepatocyte carcinoma: RCB1648), 8305C (Human thyroid-carcinoma: RCB1909), U-937 DE-4 (Human histiocytic lymphoma: RCB0435) was provided by the RIKEN Bio-Resource Center through the National Bio-Resource Project of the MEXT/AMED, Japan.

### Culture of cells

For cell culturing, a suitable medium for each cell line was used as follows: A549 – DMEM (5919; Nissui Pharmaceutical Co., Tokyo, Japan) + 10% FBS (513-96175; Corning, NY, USA); HeLa, HSC-3, HGC-27 – MEM (5902; Nissui Pharmaceutical Co., Tokyo, Japan) + 10% FBS; A431, HL 60, LNCap.FGC, U-937 DE-4, 8305 C – RPMI 1640 (5918; Nissui Pharmaceutical Co., Tokyo, Japan) + 10% FBS; HFSKF-II – HamF12 (5910; Nissui Pharmaceutical Co., Tokyo, Japan) + 15%FBS; CACO-2 – MEM + 20% FBS + 0.1 mM NEAA (11140-050; GIBCO Life Technologies, Carlsbad, CA USA); HEK293, Hep G2 – MEM + 10% FBS + 0.1 mM NEAA; MCF 7 – MEM + 10% FBS + 0.1 mM NEAA + 1 mM Sodium Pyruvate (13058-12; Nacalai tesque, INC., Kyoto, Japan); OCUB-F – DMEM + 10% FBS + 0.5 mM Sodium Pyruvate; EC-GI-10 – HamF12 + 15% FBS. The cultures were maintained and cultured at 37°C in a 5% CO^2^ incubator. Details of all the cell lines used are presented in Table 1.

**Table 1.** The Cell Lines.

| Name of Cell Line Ethnicity |  | No. | Growth properties |  |
| --- | --- | --- | --- | --- |
| HeLa | Human cervix epitheloid carcinoma | EC93021013 | Adherent | MEM + 10% FBS |
| A549 | Human alveolar adenocarcinoma | EC86012804 | Adherent | DMEM + 10% FBS |
| A431 | Human epidermoid carcinoma | EC85090402 | Adherent | RPMI1640 + 10% FBS |
| CACO-2 | Human colorectal adenocarcinoma | RCB0988 | Adherent | MEM + 20% FBS + 0.1 mM NEAA |
| HEK293 | Human embryo kidney | RCB1637 | Adherent | MEM + 10% FBS + 0.1 mM NEAA |
| HL60 | Human promyelocytic leukemia cells | RCB0041 | Floating | RPMI1640 + 10% FBS |
| HSC-3 | Human oral squamous carcinoma | RCB1975 | Adherent | MEM + 10% FBS |
| LNCap, FGC | Human prostate carcinoma | RCB2144 | Adherent | RPMI1640 + 10% FBS |
| U-937 DE-4 | Human histiocytic lymphoma | RCB0435 | Floating | RPMI1640 + 10% FBS |
| MCF7 | Human breast adenocarcinoma | RCB1904 | Adherent | MEM + 10% FBS + 0.1 mM NEAA + 1 mM Sodium Pyruvate |
| HGC-27 | Human stomach carcinoma | RCB0500 | Adherent | MEM + 10% FBS |
| OCUB-F | Human breast carcinoma | RCB0882 | Floating | DMEM + 10% FBS + 0.5 mM Sodium Pyruvate |
| EC-GI-10 | Human oesophagus carcinoma | RCB0774 | Adherent | HamF12 + 10% FBS |
| Hep G2 | Human Caucasian hepatocyte carcinoma | RCB1648 | Adherent | MEM + 10% FBS + 0.1 mM NEAA |
| 8305C | Human undifferentiated thyroid carcinoma | RCB1909 | Adherent | RPMI1640 + 10% FBS |
| HFSKF-II | Fetal skin, normal fibroblasts | RCB0698 | Adherent | HamF12 + 15% FBS |

### MTT assay

In the case of adherent cells (Table 1), 96-well culture plates (Corning Inc. New York, USA) were inoculated with 3 × 10^4^ cells / 100 μL / well of each cell and pre-cultured for 24 hours at 37 °C in 5% CO^2^. After pre-incubation, the culture was exchanged for a test medium and incubated at 37°C under 5% CO^2^ for 24 to 48 hours. To the test medium, a Gosen-Sakura leaf extract was added to a final concentration of 0.0 to 5.0% (v/v) was used as indicated in individual experiments. As a control, 0.0 to 5.0% (v/v) of PBS [(-) (Ca^2+^ / Mg^2+^ - free PBS)] (Nippon Gene Co. Ltd., Tokyo, Japan) was used.

In the case of non-adherent (floating) cells (Table 1), the culture adjusted to have 3 × 10^6^ cells / 10 mL and was pre-cultured for 24 hours at 37°C in 5% CO^2^ in a 100 mm petridish, and per 100 μL / well culture solution was dispensed in 96-well culture plates. The Gosen-Sakura cell extract was added so that the final concentration was 0.0 to 5.0% (v/v). As a control, 0.0 to 5.0% (v/v) of PBS (-) was used.

Thereafter, both adherent and non-adherent (floating) cells were cultured in the same manner. After 48 hours of incubation at 37°C in 5% CO^2^, 10 μL / well of MTT (3- [4,5-dimethylthiazol-2-yl] −0,5-diphenyl tetrazolium bromide (11465007001; Roche Life Science, Mannheim, Germany) labeling reagent (1×) was added and incubated at 37°C under 5% CO^2^ for 4 hours. Then 100 μL / well of solubilizing solution (0.01 M HCl, 10% SDS) was added, and incubated for 17 hours under the condition of 37°C under 5% CO^2^. Post-incubation, the absorbance at 570 nm was measured with an iMark microplate reader (Bio-Rad, Richmond, CA, USA) at a reference wavelength of 650 nm. The cell viability (%) in the case of addition of the test sample was determined with the number of viable cells without addition of the test sample taken as 100. Each measurement was repeated three times, and the average value was calculated.

### Component analysis of Gosen-Sakura flower and leaf extracts

Component analysis was carried out using GCMS-QP2010 SE GC/MS analyzer (Shimadzu Corporation, Kyoto, Japan). The column oven initial temperature was set at 60°C, after holding for 12 minutes, it was heated to 90°C at 3°C/min and then increased to 155°C at 5°C/min. TRACE™ TR-5 MS, 30 m × 0.25 mm ID × 0.25 μm (Thermo Fisher Scientific Inc., Waltham, MA, USA) was used as a column. Gosen-Sakura flower and leaf extract samples were standardized by injecting 1.0 μL, measured in splitless mode and quantified by the total ion count. Identification of the major compounds was done using the retention time (RT) of the GC/MS spectrum of the sample and the boiling point of the compound, MS database. For the determination of coumarin and benzaldehyde in the sample, identification was made by first dissolving coumarin and benzaldehyde in hexane and using different concentrations to prepare a calibration curve.

### Cell cycle monitoring

For cell cycle monitoring, a Cell-Clock Cell Cycle Assay Kit (Biocolor Ltd., Newtownabbey, UK) was used. Cells were seeded onto 24-well culture plates, cultured to approximately 50% confluence, replaced with fresh medium, and Gosen-Sakura leaf extract was added from 0 (blank) to 7%. For control, PBS (-), was added to same concentrations as above. The Cell-Clock dye reagent was added 24, 48 and 72 hours post-addition of Gosen-Sakura leaf extract and incubated at 37°C for 1 hour. The Cell-Clock dye reagent was removed after completion of the incubation by washing twice with the Cell-Clock dye wash reagent, and observed by a microscope (KEYENCE all-in-one fluorescence microscope BZ-X700, Osaka, Japan) within 15 minutes, and photographed. Pixel data of each color tone (yellow, green, dark blue) was acquired using ImageJ software (NIH Image, Bethesda, Maryland, USA), and the number of cells per cell cycle was calculated as a ratio from the ratio of each color tone to the total pixel value.

### Detection of apoptosis

For detection of apoptosis, APOPercentage Apoptosis Assay Kit (Biocolor Ltd., Newtownabbey, UK) was used. Cells pre-cultured under the conditions of 37°C and 5% CO^2^ were seeded on a 24-well cell culture plate at a cell density of 5 × 10^4^ cells / 500 μL / well and incubated at 37°C under 5% CO^2^ conditions for 24 hours. Gosen-Sakura leaf extract was added at 0 to 7%, and PBS (-) was added to the negative control. Thereafter, culturing was carried out at 37°C and 5% CO^2^ for 6, 18, and 24 hours. For positive control of apoptosis, 1 μL of H_2_O_2_ (Wako, Tokyo, Japan) was added per 500 μL / well and cultured for 2 hours at 37°C and 5% CO^2^. After culturing, the culture medium was withdrawn from each well, and 500 μL of a medium containing 5% APOPercentage Dye was added followed by incubation at 37°C under 5% CO^2^ for 30 minutes. Post-incubation, the solution was withdrawn from each well, and gently washed twice with 1000 μL of PBS (-) to remove the APOPercentage Dye. Thereafter, 500 μL PBS was added and observed with a microscope (KEYENCE all-in-one fluorescence microscope BZ-X700, Osaka, Japan) and images were taken.

### DAPI staining

The cells were pre-cultured at 37°C under 5% CO^2^ condition, the cells were plated so that 5 × 10^4^cells / 1000 μL / well in 4-well cell culture plates (Iwaki Asahi Glass Co. Ltd., Tokyo, Japan), 37°C, for 24 hours under 5% CO^2^ condition. After replacing with fresh medium, Gosen-Sakura leaf extract was added at concentration of 0 ∼ 7%; PBS (-) was added as a negative control. Thereafter, culturing was carried out at 37°C and 5% CO^2^ at time points for 18 and 24 hours. To the positive control, 2 μL of H_2_O_2_ was added per 1000 μL / well and cultured under the condition of 37°C and 5% CO^2^ for 2 hours. Post-incubation, 500 μL of acetone which had been cooled by washing with PBS (-) was added and kept at −0°C for 10 minutes. Thereafter, washing was carried out three times with PBS (-), 1000 μL of 0.1% DAPI (D 1306; Thermo Fisher Scientific, Waltham, MA, USA) / PBS (-) was added and incubated at 37°C under 5% CO^2^ for 20 minutes. Finally, washing was carried out three times with PBS (-), and the image was taken by observation with a microscope (KEYENCE all-in-one fluorescence microscope BZ-X700, Osaka, Japan). OP-87762 BZX Filter was used for photographing for DAPI (excitation wavelength 360/40, absorption wavelength 460/50).

## Results and discussion

### Low temperature extraction method for the Gosen-Sakura leaves

The harvested Gosen-Sakura (Gosen-city, Niigata prefecture) [15] petals and leaves were transported at room temperature to the low-temperature vacuum facility and processed (cold vacuum extraction) to obtain the flowers and leaves extract (and cell dry powder) (Fig 1). The low-temperature vacuum extraction method is a relatively inexpensive technology developed recently in Japan (with an International Patent; [16]) and with the greatest merit that the extraction occurs in a nearly natural form due to the process being carried out at a low temperature of around 40-45°C (Fig 1; see also [17]). Briefly, the essential oil and floral (or tissue) water are separated by the steam distillation method, but ‘cell extract’ is extracted by low temperature vacuum extraction method. When extracting essential oil by steam distillation method, floral water is a mixture composed of trace amounts of essential oil components in steam obtained as a byproduct, while the ‘cell extract’ is water-soluble as it contains the plant moisture content. Because there are many essential oil components and the content of plants components is high, the cell extract can be said to be a ‘water-soluble aroma’. As solvents and water do not mix, the cell extracts are highly safe (non-toxic) and can be used as raw materials in the cosmetics and health food industry. It has been confirmed that the aroma components from the residues of Okinawa Shikuwasa (http://www.jpn-okinawa.com/en/products/shiikwaasaa/) juice using low temperature vacuum extraction are about 10 times to 100 times as large as the essential oil obtained by the steam distillation method. Moreover, 100% natural flavor ingredients can be extracted from plants such as “strawberries” and “cherry blossoms” which had been previously considered impossible to extract.

The low temperature vacuum extraction method has the following features: i) since extraction is performed at a low temperature of around 40°C, components sensitive to heat can be extracted in a state close to natural, ii) since solvents, steam, etc., are not used at all, it is possible to extract aromatic substances derived from the 100% raw materials, and, iii) the technology is based on an applied machine being commercialized as a dryer, and it is relatively inexpensive and easy to operate.

In the case of the Gosen-Sakura, following the low temperature vacuum extraction process, we obtained the leaf extract (around 95%) and cell dry powder (5%). These were used for downstream experiments as described below.

### Study on growth inhibitory effect of Gosen-Sakura leaf extract on human tumor cell line

The results confirming the proliferation inhibitory effect of Gosen-Sakura leaf extract on 15 tumor cells by MTT assay is shown in Fig 2. Among the 15 human tumor cells tested, those with particularly high sensitivity to the Gosen-Sakura leaf extracts are HeLa (cervical cancer), A431 (epithelial-like cell cancer), HL60 (promyelocytic leukemia cell), U-937 DE-4 (blast cell leukemia), HGC-27 (gastric cancer), and Hep G2 (liver cancer). When 5% of the Gosen-Sakura leaf extracts was added, the cell survival rate was suppressed to 20% or less. The next most sensitive cell lines were HSC-3 (tongue squamous cell carcinoma) and EC-GI-10 (esophageal squamous cell carcinoma), and the cell viability increased from 20% to 30% under 5% Gosen-Sakura leaf extracts. The effect was less comparable on the remaining 7 strains of A549 (lung cancer), CACO-2 (colon cancer), HEK293 (derived from fetal kidney cells), LNCap.FGC (prostate cancer), MCF7 (breast cancer), OCUB-F (breast cancer), 8305C (thyroid cancer); i.e., the sensitivity was low as compared with the above strains.

**Figure 2.**
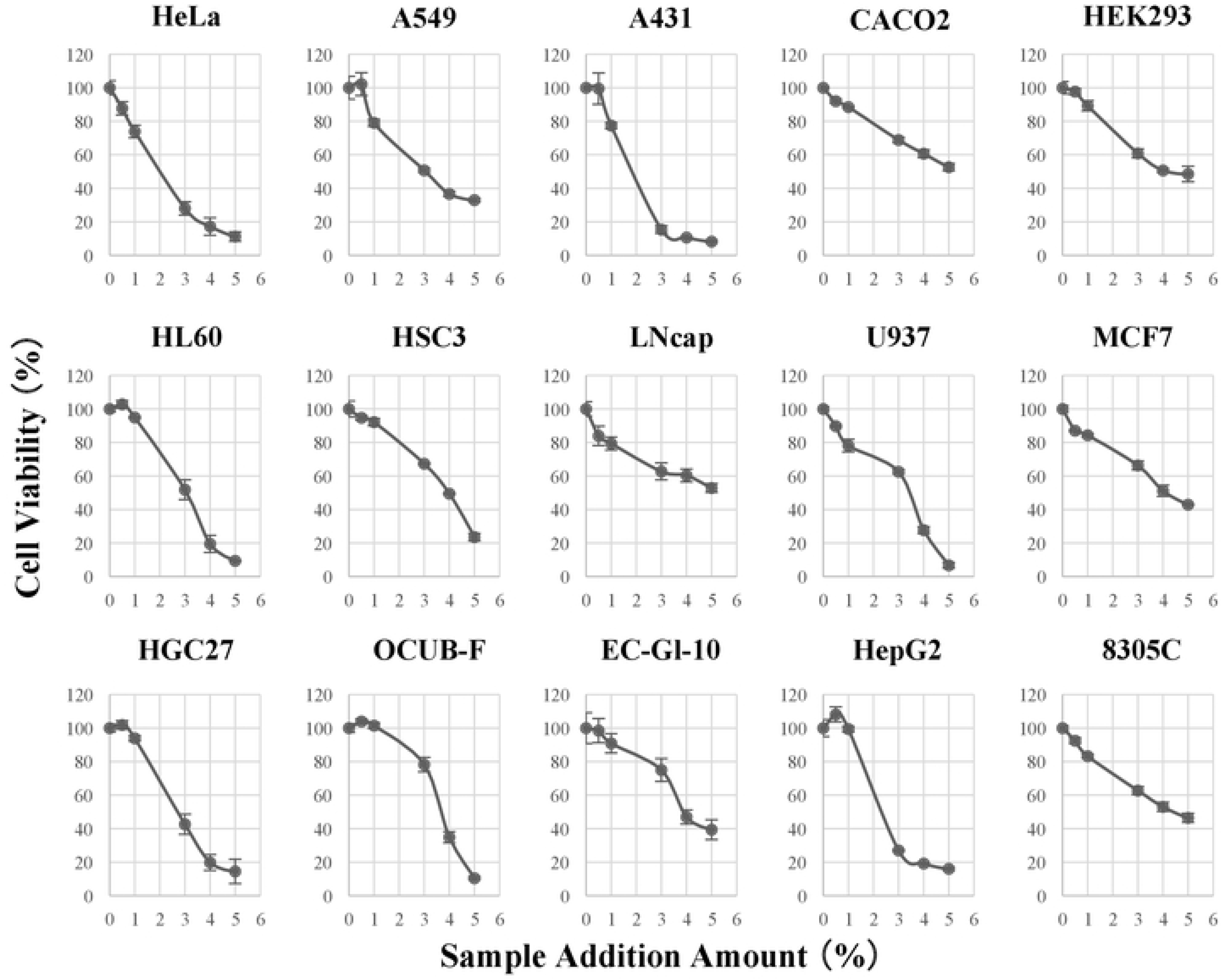
Inhibitory effect of the double cherry blossom leaf extracts on the proliferation 15 tumor cells lines by MTT assay. The cell line details are presented in Table 1.

The results confirming the growth inhibitory effect of Gosen-Sakura leaf extracts on normal cells is shown in Fig 3. Using the MTT assay, we examined the effect of Gosen-Sakura leaf extract in normal cells, at 6, 18 and 24 hours post-treatment. In the case of a normal cell, HFSKF-II (normal fetal skin-derived fibroblasts) was used and the growth inhibitory effect by Gosen-Sakura leaf extract was confirmed. From the results shown in Fig 3, it was confirmed that the proliferation inhibitory effect by Gosen-Sakura leaf extract was higher when the treatment time was longer up to 24 hours in the case of both normal cells and tumor cells. However, in the case of HeLa cells (tumor cell line), the proliferation inhibition rate was significantly much higher as compared to the normal cells (HFSKF-II) under Gosen-Sakura leaf extract exposure from 3% up to 7%. Based on the results of this experiment, it can be suggested that the susceptibility to Gosen-Sakura leaf extract changes depending on the exposure time and sensitivity to Gosen-Sakura leaf extract may be low depending on the strain.

**Figure 3.**
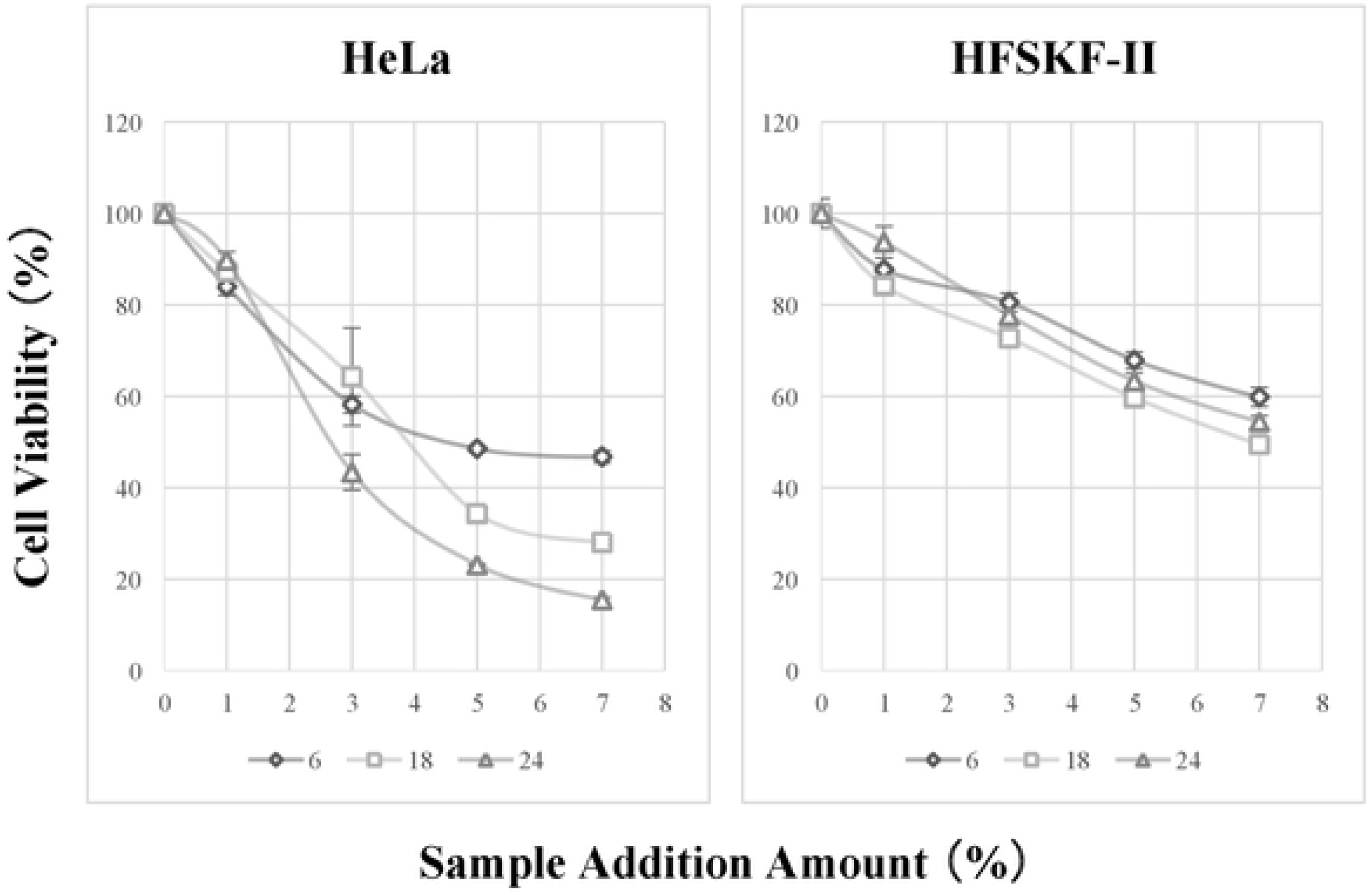
Confirming the growth inhibitory effect of double cherry blossom leaf extracts on normal cell (HFSKF-II) with HeLa cell.

### Component analysis results of the Gosen-Sakura leaf extract

We detected coumarin and benzyl alcohol as main components of the Gosen-Sakura leaf extract; and, it was confirmed that coumarin accounted for 70% or more and benzyl alcohol for around 20%. Quantification of these 2 components revealed coumarin to be 75 μg / 1000 mL (molar concentration 500 nM) and 20 μg / 1000 mL of benzyl alcohol; we could not quantity the molar concentration of benzyl alcohol in this study. Therefore, based on the results of the component analysis, we confirmed the antitumor effect on HeLa cell line at concentrations of 75 μg / 1000 mL of coumarin and 20 μg / 1000 mL of benzyl alcohol, but the proliferation inhibitory effect similar to that of the Gosen-Sakura leaf extract was not observed. However, combining benzyl alcohol with coumarin, a growth inhibitory effect was observed. However, since the cell viability did not decrease below 60% or less even under the added condition of 5% (benzyl alcohol with coumarin), it was suggested that the Gosen-Sakura leaf extract contains a substance having antitumor effect in addition to the main components coumarin and benzyl alcohol (Fig 4).

**Figure 4.**
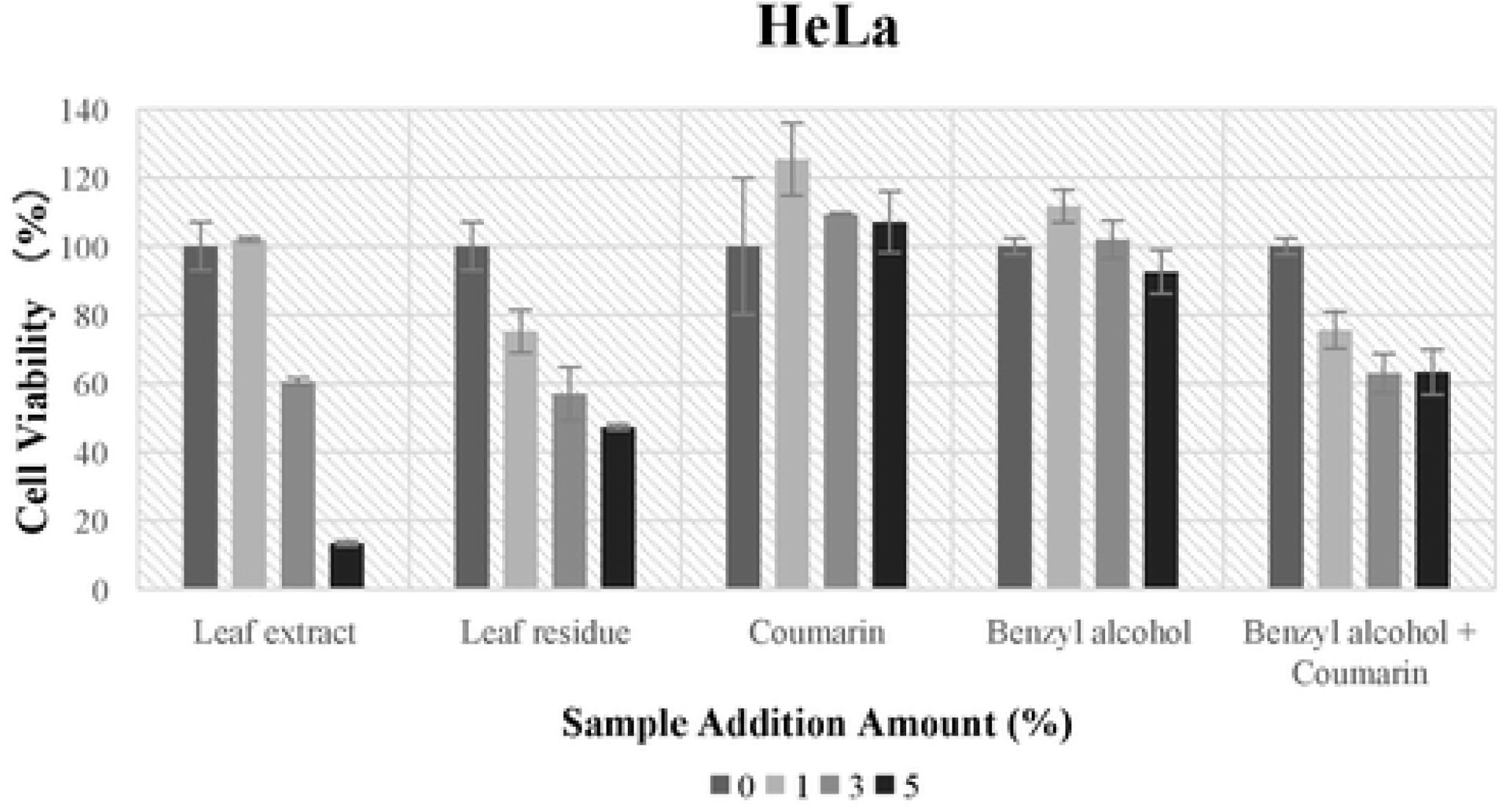
Double cherry blossom leaf extract contains a substance having antitumor effect, in addition to the main components coumarin and benzyl alcohol, as shown by its effects on HeLa cells.

### Effect of Gosen-Sakura leaf extract on the cell cycle

Using the Cell-Clock Cell Cycle Assay Kit we examined the staining of HeLa and A549 cells after treatment with Gosen-Sakura leaf extract for 48 hours. Each color tone (G1 stage: yellow, S phase: green, G2 And M phase: dark blue) and the number of cells per cell cycle was calculated as a ratio from the ratio of each color tone to the total pixel value as shown in Fig 5. The cell cycle is divided into M phase and interphase (G1 phase, S phase, G2 phase). Furthermore, cells that have temporarily or reversibly discontinued division comprise the G0 phase. In each period, M phase is the cell division phase, G1 phase is geared for DNA synthesis, S phase is the DNA replication (DNA synthesis) phase, and the G2 phase is geared for mitosis and cytokinesis (M phase). From the results in Fig 5, it was confirmed that for HeLa cells G1 phase was observed at 48 hours and S phase was increased at 72 hours. For A549 cells, the G1 phase tended to increase at both 48 h and 72 h. Based on the results of this study, it was confirmed that the proportion of both cells lines showing G1 phase and S phase increased by stagnation in G1 phase and S phase following treatment with the Gosen-Sakura leaf extract.

**Figure 5.**
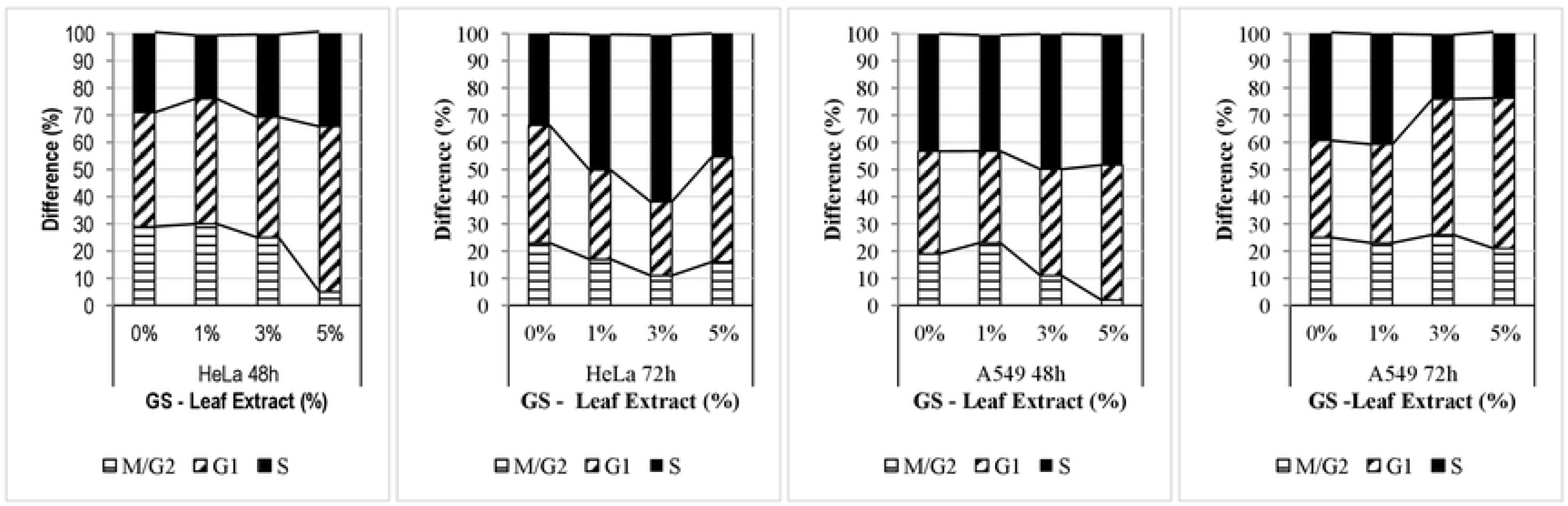
Staining of the HeLa and A549 cells by the Cell-Clock Cell Cycle Assay Kit after treatment with the double cherry blossom leaf extract for 48 housing. Each color tone (G1 stage: yellow, S phase: green, G2 And M phase: dark blue) and the number of cells per cell cycle was calculated as a ratio from the ratio of each color tone to the total pixel value.

Since the G1 and S phases are the preparation period for DNA synthesis and the period of DNA replication (DNA synthesis) as described above, it is suggested that the tumor cell proliferation inhibitory action by the double cherry blossom leaf extract is during the process of DNA synthesis. Furthermore, it can be considered that the double cherry blossom leaf extract causes apoptosis because it has been clarified that when apoptosis occurs during the DNA repair mechanism, DNA damage cannot completely repaired. Therein, the rationale for the next experiment as performed below.

### Gosen-Sakura leaf extract induces apoptosis

Results of the HeLa (Fig 6) and A549 (Fig 7) tumor cells by APOPercentage Apoptosis Assay Kit and DAPI staining reveal the effects of Gosen-Sakura leaf extract as also suggested from the above results. The upper panel of each Figs 6 and 7 shows the stained image by APOPercentage Apoptosis Assay Kit, the lower panel shows the stained image by DAPI. The left panels in each Figs 6 and 7 show the cells treated with H_2_O_2_ (hydrogen peroxide) as a positive control, and as a negative control the Gosen-Sakura leaf extract without addition (0%) and with 3%, 5% and 7% Gosen-Sakura leaf extract, respectively. The results confirmed that the number of cells stained red (upper panels), that is, the number of apoptosis-induced cells increased with an increase in the amount of the Gosen-Sakura leaf extract. Almost all tumor cells showed apoptosis under the conditions of 7% cherry leaf cell extract treatment for 18 hours. Further, the DAPI stained images (lower panels) confirmed that increase in the amount of the Gosen-Sakura leaf extract, the nuclei were condensed; i.e., they were small. Since it is known that this nuclear aggregation phenomenon is a morphological change peculiar to apoptosis, apoptosis induction of HeLa cells is thought to be result of the Gosen-Sakura leaf extract. Similarly, for the A549 cells, images with the Apoptosis Assay Kit confirmed that the number of cells stained red staining became larger with an increase in the Gosen-Sakura leaf extract treated samples. However, unlike HeLa cells, the whole cells did not turn red, but condensation of the nucleus was confirmed by DAPI staining, so the possibility of apoptosis induction by the Gosen-Sakura leaf extract on A549 cells could be highly considered.

**Figure 6.**
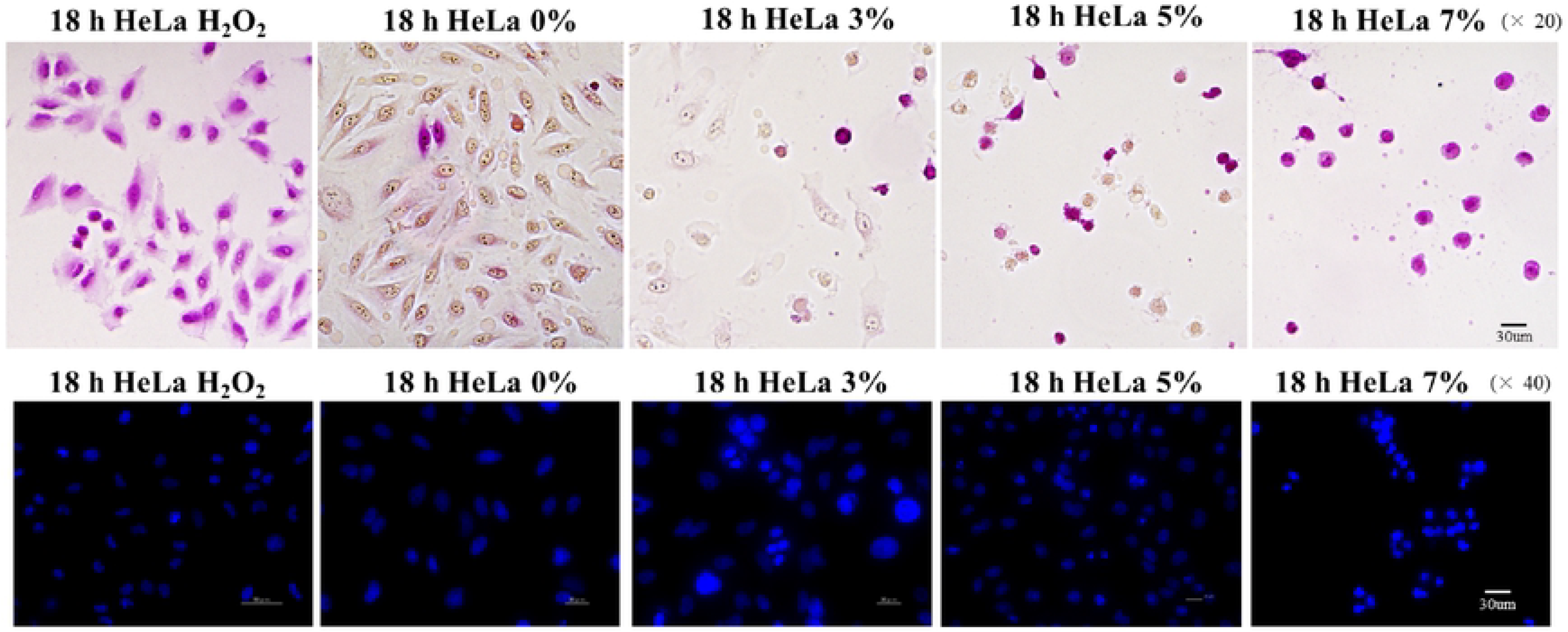
APOPercentage Apoptosis Assay Kit and DAPI staining of HeLa tumor cells.

**Figure 7.**
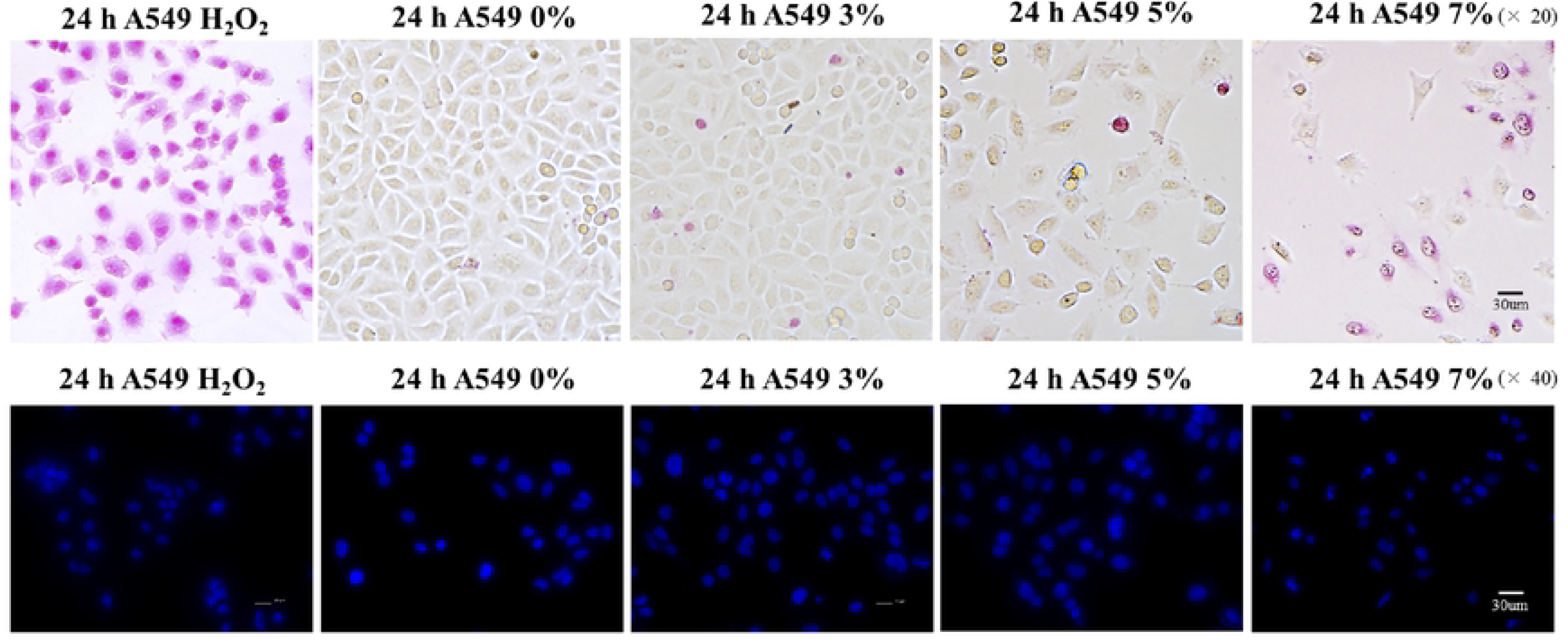
APOPercentage Apoptosis Assay Kit and DAPI staining of A549 tumor cells.

### Concluding remarks

These results demonstrate the presence of ‘novel’ bioactive compound(s) in the nearly natural form of a water-soluble leaf (cell) extract of the Gosen-Sakura leaves. The novelty of the study is once again highlighted in the fact that the Gosen-Sakura leaves extraction was carried out using low temperature and where the method was established for the leaves for the first time; previously we the group of Shioda and co-workers had reported the protocol for the lavender flower oil [17]. Secondly, the water-soluble nature of the Gosen-Sakura leaf extracts suggests the possibility of not only polyphenols but also other compounds that have not yet been identified. This research also demonstrated the proliferation inhibitory effect of Gosen-Sakura leaf extract on multiple tumor cells by the MTT assay. To our knowledge, no reports exist on the sakura tree ‘leaf (cell) extracts inhibiting tumor cell line growth. However, organic (methanolic) solvent extracts of cherry blossoms showed not only high phenol content, radical scavenging activity and reducing power, but also inhibition of the growth of human colon cancer cell line HT-29 [6]. Moreover, as the cherry leaves contain coumarin, it was examined how the Gosen-Sakura leaf extract compared with coumarin and other known compounds such as benzyl alcohol for anti-tumor affects. As the combination of benzyl alcohol with coumarin did not show a high-level of growth inhibitory effect we hypothesize that the Gosen-Sakura leaf extract contains another substance(s) with antitumor effect. Our research further implicated the anti-tumor action (tumor cell proliferation inhibition) of Gosen-Sakura leaf extracts occurred during the process of DNA synthesis. With this rationale, we examined the apoptosis in tumor cells, and clearly demonstrated the occurrence of apoptosis under the conditions of 7% cherry leaf cell extract treatment for 18 hours in almost all cell lines. The next step of the research will attempt the identification (through high-performance liquid chromatography separation in conjunction with tandem mass spectrometry) of this potential ‘novel’ anti-tumor compound(s) from the Gosen-Sakura leaves.

## Acknowledgments

This study was supported in part by Gosen-City in Niigata Prefecture, Japan and also by the Ministry of Agriculture, Forestry and Fisheries of Japan.

